# Cardiac Metabolic Limitations Contribute to Diminished Performance of the Heart in Aging

**DOI:** 10.1101/560649

**Authors:** X. Gao, D. G. Jakovljevic, D. A. Beard

## Abstract

Changes in the myocardial energetics associated with aging—reductions in creatine phosphate (CrP)/ATP ratio, total creatine, and ATP—mirror changes observed in failing hearts compared to healthy controls. Similarly, both aging and heart failure are associated with significant reductions in cardiac performance and maximal left ventricular cardiac power output (CPO) compared to young healthy individuals. Based on these observations, we hypothesize that reductions in the concentrations cytoplasmic adenine nucleotide, creatine, and phosphate pools that occur with aging impair the myocardial capacity to synthesize ATP at physiological free energy levels, and that the resulting changes to myocardial energetic status impair the mechanical pumping ability of the heart. The purpose of this study is to test theses hypotheses using an age-structured population model for myocardial metabolism in the adult female population and to determine the potential impact of reductions in key myocardial metabolite pools in causing metabolic/energetic and cardiac mechanical dysfunction associated with aging. To test these hypotheses, we developed a population model for myocardial energetics to predict myocardial ATP, ADP, CrP, creatine, and inorganic phosphate concentrations as functions of cardiac work and age in the adult female population. Model predictions support our hypotheses and are consistent with previous experimental observations. The major findings provide a novel theoretical and computational framework for further probing complex relationships between the energetics and performance of the heart with aging.

**Significance:** Normal mechanical function of the heart requires that ATP be continuously synthesized at a hydrolysis potential of roughly −60 kJ mol^−1^. Yet in both the aging and diseased heart the relationships between cardiac work rate and concentrations of ATP, ADP, and inorganic phosphate are altered. Important outstanding questions are: To what extent do changes in metabolite concentrations that occur in aging and heart disease affect metabolic/molecular processes in the myocardium? How are systolic and diastolic functions affected by changes in metabolite concentrations? This study addresses these questions by analyzing relationships between cardiac energy demand and supply using an age-structured population model for human myocardial energetics in women.

## Introduction

Peak left ventricular cardiac power output (CPO), as an integrative measure of overall pumping capability and performance of the heart, is the strongest predictor of mortality in heart failure [1]. Yet even in the absence of heart failure, maximal CPO decreases with age and thus increases the risk cardiovascular morbidity and mortality [2]. Changes in myocardial energetics with age are associated with diminished maximal CPO in older people [3, 4]. These observations raise the questions of if and how age-dependent changes to myocardial metabolic energy supply influence the myocardial capacity to generate power. Previous studies have established a theoretical link between reductions in cytoplasmic adenine nucleotide, creatine, and phosphate metabolite pools and altered myocardial energetics in heart failure [5]. The goals of the present study are to determine whether or not reductions in these metabolite pools can potentially explain age-dependent changes in myocardial energetics in normal subjects and whether or not the resulting energetic changes can potentially explain the observed reductions in maximal CPO with age.

Left ventricular CPO is the rate at which the ventricle produces mechanical work to pump blood through the systemic circulation. CPO is calculated as left ventricular stroke work (mechanical work done per heart beat) multiplied by the heart rate, or as average difference in pressure between the left atrium and the aorta multiplied by the cardiac output, expressed in units of energy per unit time. Processes such as cross-bridge cycling require that ATP is continuously synthesized (and ADP and inorganic phosphate consumed) at sufficient concentrations such that normal functions are not kinetically or thermodynamically impaired [6, 7]. Because demand for ATP production in the myocardium is proportional to cardiac work, and because ATP hydrolysis serves as the source of chemical free energy to drive myocardial contraction, it stands to reason that limitations or impairments in myocardial energy metabolism may influence overall cardiac performance represented by CPO.

It is observed that in both the aging [3, 4] and the diseased heart [8–12] relationships between cardiac work rate and concentrations of phosphate metabolites ATP, ADP, and phosphocreatine (CrP) are altered. In heart failure ATP and the CrP/ATP ratio is diminished and [ATP] in myocardium is lower compared to healthy individuals [8, 12–14]. Although heart failure is a complex syndrome which can arise due to a variety of pathophysiological abnormalities [15], alterations to the myocardial energetics state are established hallmarks of heart failure irrespective of etiology [8, 13, 16, 17]. While a complete understanding of the mechanistic links between energy metabolism and mechanical function is lacking, energy metabolism represents an actively pursued potential target for treating heart failure [18–21].

Similarities between the myocardial energetic phenotypes observed in heart failure/decompensation and aging include lower myocardial ATP and CrP/ATP compared to the healthy young heart [4, 22]. Based on these similarities and similarities in terms of diminished maximum cardiac performance, we hypothesize that reductions in the concentrations cytoplasmic adenine nucleotide, creatine, and phosphate pools that occur with aging impair the myocardial capacity to synthesize ATP at physiological free energy levels, and that the resulting changes to myocardial energetic status play a causal role in contributing to reductions in cardiac performance with aging.

To explore the viability of these hypotheses we applied a model of myocardial energetics previously used to analyze data from large animals [5, 23]. An age-structured population model for myocardial metabolite pools was parameterized based on studies in humans from Kostler et al. [22] and Jakovljevic et al. [3] to represent variability and interrelationships between adenine nucleotide, creatine, and phosphate pools in the human female population. The model of population variability in metabolite pools was used in conjunction with simulations of myocardial energetics to predict the relationship between maximum CPO and age. Because of a significant difference in age-associated changes in cardiac morphology and function between men and women [2, 24, 25], the present model accounts for female subjects only. Model predictions are consistent with observations on the relationships between age and maximum CPO in female subjects [3, 4] and consistent with our hypotheses. These simulations provide a theoretical/computational framework for further probing relationships between energetic status and cardiac power and for comparing predictions associated with the hypothesis and experimental data.

## Methods

### Overview of simulation methods

A previously developed model of myocardial energetics developed by Wu et al. [5, 23] was used to simulate how myocardial oxygen consumption 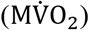 and cellular concentrations of ATP, ADP, inorganic phosphate (Pi), Cr, and CrP vary with changes in ATP demand. This model has been validated on the basis of its ability to match steady-state data on how these variables vary with changes in ATP demand under physiological conditions, how CrP and Pi levels change dynamically in response to acute ischemia and recovery, and how these variables are altered in response to changes in concentrations of key cytoplasmic metabolite pools. Using this underlying model framework, an age-structured population model for the total adenine nucleotide (TAN), total creatine (CRtot), and total exchangeable phosphate (TEP) metabolite pools is parameterized from data on resting myocardial CrP/ATP and [ATP] in healthy women aged 20-81 years [3, 22]. The relationship between 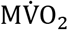 and CPO is determined empirically based on observations showing a linear relationship between oxygen consumption and power output over a broad range of physiological conditions in animal models and man.

### Population myocardial energetics model

The model of Wu et al. [5, 23] was used to simulate myocardial oxygen transport, mitochondrial tricarboxylic acid cycle kinetics and oxidative phosphorylation, and cytoplasmic adenylate kinase, creatine kinase, and ATP hydrolysis. Model predictions of how phosphate metabolite levels vary with 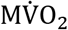 are illustrated in Figure 1 for 20 year-old subjects. To obtain these results the ATP demand (the cellular ATP hydrolysis rate) is varied from 0.36 to 1.52 mmols^−1^(1 cell)^−1^. This range of ATP hydrolysis rate yields a range of 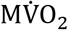 of 3.5 to 13.5 mmol·min^−1^·g^−1^ (or 78 to 302 μlO_2_·min^−1^·*g*^−1^). Since 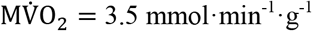 is the mean value of LV myocardial oxygen consumption observed in anesthetized dogs under baseline conditions in the data sets analyzed by Wu et al. [23] and 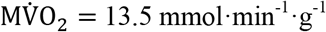 represents the upper limit of LV oxygen consumption in conscious exercising dogs [26], we use this range of ATP demand to capture upper and lower limits of the physiological range. Simulations summarized in Figure 1 show that mean cytoplasmic inorganic phosphate concentration ([Pi]_*c*_) increases from 0.43 mM under resting conditions (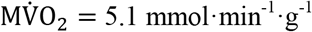 [27]) to 2.0 mM at a moderate exercise intensity 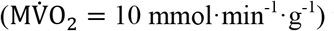 to 4.3 mM at maximal exercise conditions 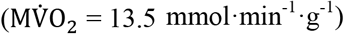, while the CrP/ATP ratio decreases from approximately 2.06 at rest to 1.80 at 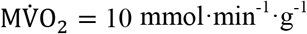 and to 1.50 at maximal exercise. These concentration ranges correspond to the expected ranges based on available in vivo data [23, 28].

**Figure 1.**
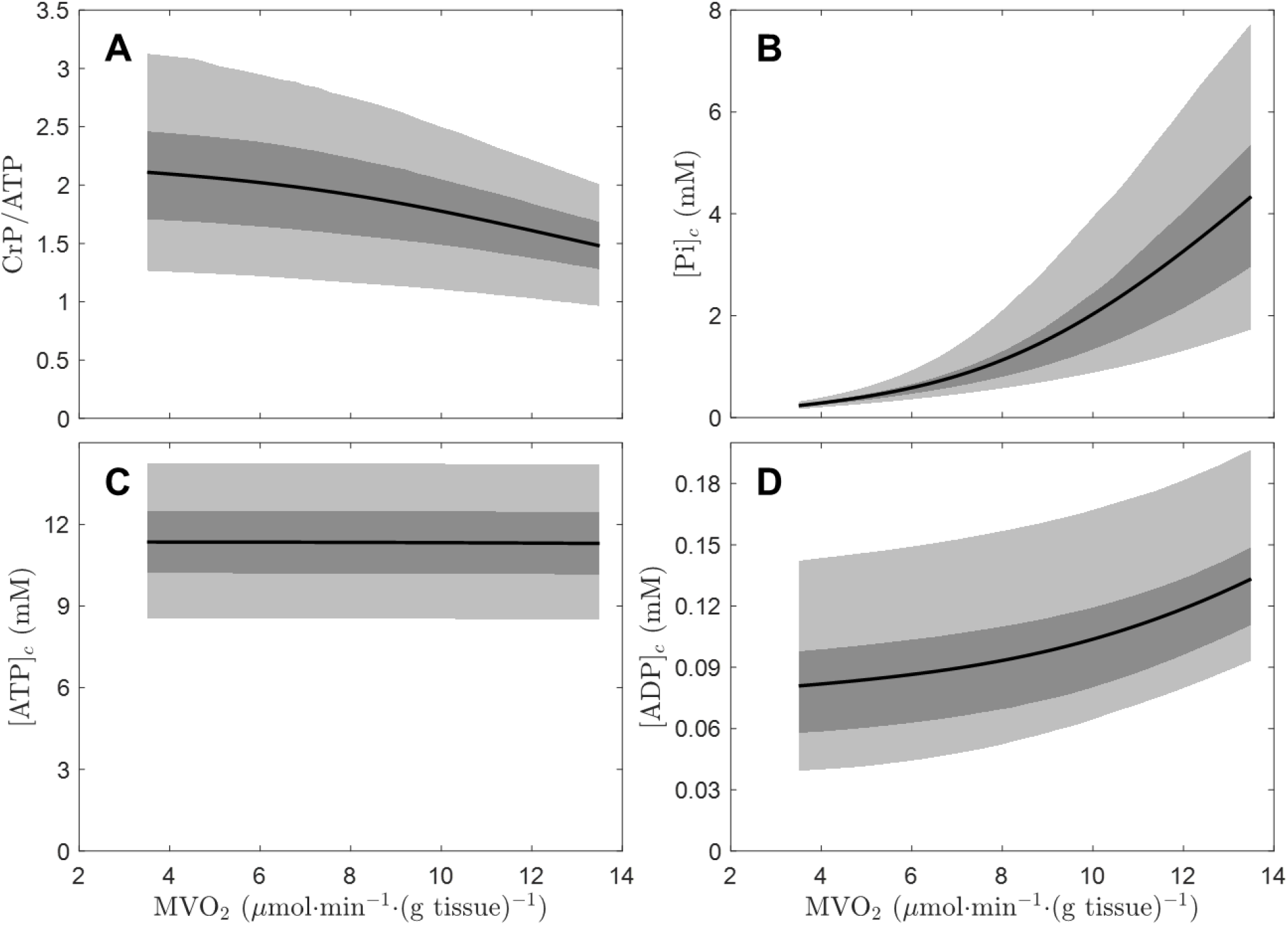
Myocardial cellular energetics. Population model predictions of myocardial energy metabolite levels are plotted as functions of 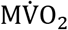. Model-predicted CrP/ATP, cytoplasmic inorganic phosphate concentration [Pi]_*c*_, cytoplasmic ATP concentration [ATP]_*c*_, and cytoplasmic ADP concentration [ADP]_*c*_ are plotted for a population of 20 year-old female subjects. The light shaded areas represent the 90% confidence range from the population prediction for each variable; the dark shaded areas represent the 50% confidence range; and the solid line represents population mean.

Predictions of the Wu et al. model depend on the levels of the three conserved metabolite pools: the total adenine nucleotide pool (TAN), the total exchangeable phosphate pool (TEP), and the total creatine pool (CR_tot_). These pools are defined as:

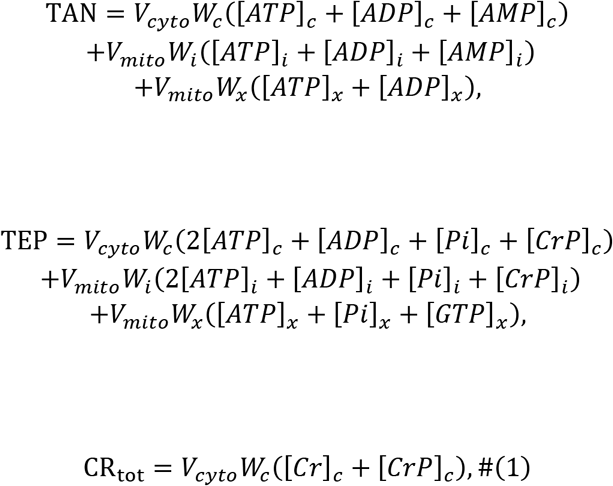

where *V_cyto_* and *V_mito_* are the cellular volume fraction of cytoplasm and mitochondria, respectively; *W_c_, W_i_*, and *W_x_* are water volume fraction in cytoplasm, mitochondrial intermembrane space, and mitochondrial matrix, respectively; solute concentration in different cellular region is represented by the solute name in a square bracket together with a subscript “c”, “i”, or “x” indicating cytoplasm, mitochondrial intermembrane space, and mitochondrial matrix, respectively. The values of volume fractions used in this study are from Wu et al.

It is assumed that the mean level of each metabolite pool varies linearly with human age, and the distribution of pool level at each specific age follows the normal distribution. Thus, for an individual subject, the metabolite pool is defined with a linear model as:

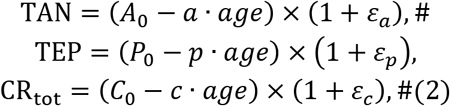

where *A*_0_, *P*_0_, *C*_0_, *a, p*, and *c* are parameters, and *ε_a_, ε_p_*, and *ε_c_* are random variables assumed to follow normal distributions with a mean of zero and variances defined

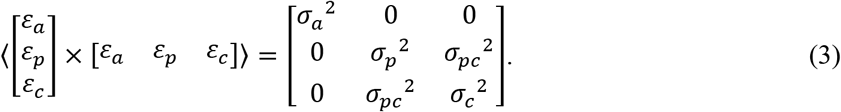

Equation (3) assumes that the variability in the phosphate and creatine pools are correlated (with correlation coefficient *σ_pc_*) while the adenine nucleotide pool varies independently of the other pools.

Values of parameters *A*_0_, *P*_0_, *C*_0_, *a, p, c, σ_a_, σ_p_, σ_c_*, and *σ_pc_* are listed in Table 1. Values of these parameters were estimated as follows. The mean levels for all three metabolite pools in the 20-year-old population were assumed to be equal to the normal levels determined by Wu et al. [5]:

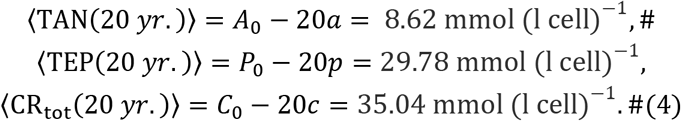

**Table 1:** Population model parameters

|  | Definition | Value | Units |
| --- | --- | --- | --- |
| $A_0$ | Reference TAN value, assigned from Equation (4) | 8.62 | $\text{mmol (l cell)}^{-1}$ |
| $P_0$ | Reference TEP value, assigned from Equation (4) | 29.78 | $\text{mmol (l cell)}^{-1}$ |
| $C_0$ | Reference $\text{CR}_{\text{tot}}$ value, assigned from Equation (4) | 35.04 | $\text{mmol (l cell)}^{-1}$ |
| $a$ | Slope of relationship between mean TAN and age, estimated by fitting data in Fig. 2 | 0.082 | $\text{mmol (l cell}\cdot\text{yr.)}^{-1}$ |
| $p$ | Slope of relationship between mean TEP and age, determined by Equation (5) | 0.283 | $\text{mmol (l cell}\cdot\text{yr.)}^{-1}$ |
| $c$ | Slope of relationship between mean $\text{CR}_{\text{tot}}$ and age, estimated by fitting data in Fig. 2 | 0.39837 | $\text{mmol (l cell}\cdot\text{yr.)}^{-1}$ |
| $\sigma_a$ | Relative variance in TAN pool, estimated by fitting data in Fig. 2 | 0.12 | unitless |
| $\sigma_p$ | Relative variance in TEP pool, estimated by fitting data in Fig. 2 | 0.06 | unitless |
| $\sigma_c$ | Relative variance in $\text{CR}_{\text{tot}}$ pool, estimated by fitting data in Fig. 2 | 0.12 | unitless |
| $\sigma_{pc}$ | Covariance between TEP and $\text{CR}_{\text{tot}}$ pool, estimated by fitting data in Fig. 2 | 0.08 | unitless |

Furthermore, the relationship between *a* and *p* was fixed to maintain the average proportion between TAN and TEP determined by Wu et al.:

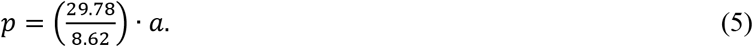

Using the relationships defined in Equations (4) and (5), values of *A*_0_, *P*_0_, *C*_0_, and *p* and calculated from estimates of *a* and *c*. The six adjustable parameters (*a, c, σ_a_, σ_p_, σ_c_*, and *σ_pc_*) defining the age-dependent variability in the metabolite pools are estimated by fitting model predictions to data from Köstler et al. [22] and Jakovljevic et al. [3], as detailed below.

Population model results were obtained by randomly sampling the distributions of TAN, TEP, and CR_tot_ governed by Equations (2) and (3). Results below were obtained by simulating 13 fixed-age groups representing ages from 20 to 80 years. Within each age group 1000 independently determined individuals were simulated. Simulations representing the resting state are obtained setting the cellular ATP hydrolysis rate to 0.547 mmol·s^−1^·(1 cell)^−1^, which yields a resting state 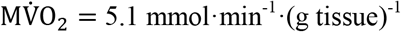.

## Results

### Parameterization of population model

Model predictions for resting-state myocardial energetics are compared to data from Köstler et al. [22], Jakovljevic et al. [3] in Figure 2. The mean model outputs are plotted a solid black lines. The light shaded areas represent the 90% confidence range from the population prediction for each variable and the dark shaded areas represent the 50% confidence range.

**Figure 2.**
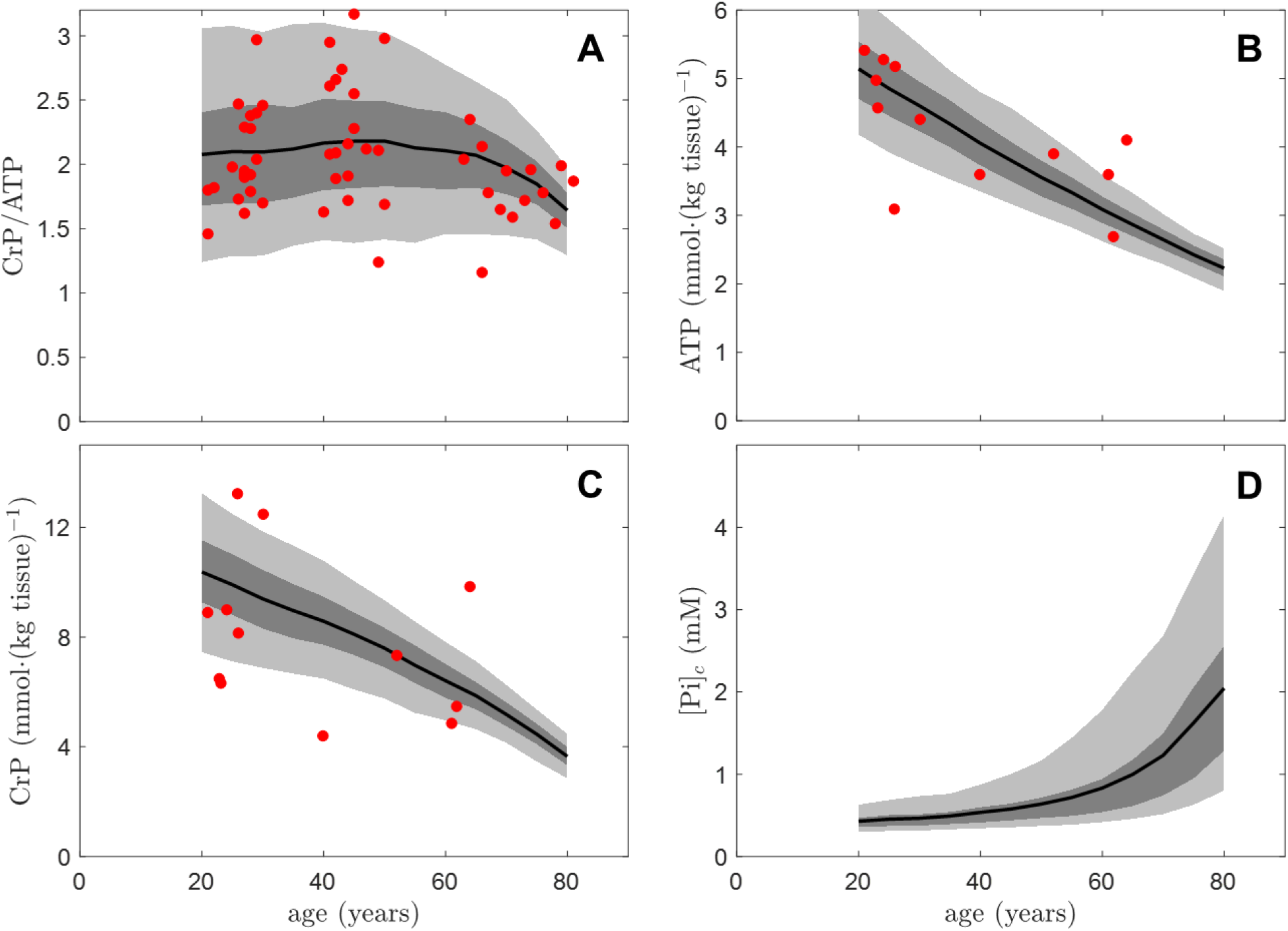
Changes in resting-state myocardial cellular energetics with age. Population model predictions of resting-state myocardial energy metabolite levels are plotted as functions age. Model-predicted CrP/ATP (panel A), total ATP (panel B), total CrP (panel C), and cytoplasmic inorganic phosphate concentration [Pi]_*c*_ (panel D), are compared to data from Köstler et al. [22] and Jakovljevic et al. [3]. The light shaded areas represent the 90% confidence range from the population prediction for each variable; the dark shaded areas represent the 50% confidence range; and the solid line represents population mean. Data from individual subjects are plotted as red dots.

Figure 2A compares results on resting CrP to ATP ratio to data from Jakovljevic et al. [3] obtained from 55 healthy untrained women with age ranging from 21 to 81 years. The population model, defined based on Equations (2) and (3) effectively captures the trends observed by Jakovljevic et al. Data from 29 of the 55 individuals fall within the 50% confidence range. Data from 3 of the 55 individual fall outside of the 90% confidence range. Panels B and C of Figure 2 compare results on resting myocardial ATP and CrP levels to data from Köstler et al. [22] from 12 healthy women ranging from 21 to 64 years old. In panels B and C, simulations and data are reported in terms of total ATP and CrP per mass of myocardium. Panel D plots the associated model predictions to cytoplasmic inorganic phosphate concentration [Pi]_*c*_, which is reported here in units of millimoles per liter of myocyte cytoplasmic water space. The predicted decreases in CrP and ATP and with increasing age are in good agreement with the trends in the data from Köstler et al.

The population model predicts that resting-state cytoplasmic [Pi] increases from approximately 0.5 mM in 20-40 year-old individuals to 1.0 mM in the mean 65 year-old, and to greater than 1 mM in most individuals aged 70 years and older.

### Prediction of maximum cardiac power output

To predict maximal CPO the maximum 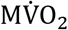 was estimated for each individual in the virtual populations by increasing cytoplasmic ATP demand until the cytoplasmic inorganic phosphate concentration reached 4.4 mM, the value associated with the maximal 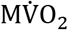 in the mean 30 year-old individual. The maximal 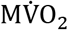 was converted to maximal CPO based on the proportionality between 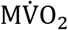 and work:

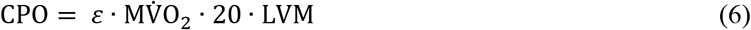

where the myocardial external efficiency *ε* is set to 0.25 [29], 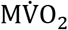 is expressed in units of mL·(second g tissue)^−1^, the caloric equivalent of 1 mL of O_2_ is taken to be 20 Joules, and LVM is the LV mass set to 150 g.

Figure 3 compares model-predicted maximum CPO to data from Jakovljevic et al. [3]. Both panels plot the model predictions as in Figures 1 and 2 with the light shaded areas represent the 90% confidence range from the population prediction for each variable and the dark shaded areas represent the 50% confidence range. The left panel plots the individual data points as red dots. The right panel plots statistical summaries of the data for three age groups, showing mean age and mean maximum CPO, and with error bars indicating ± 1 standard deviation. The model predicts that from age 20 to 80, the mean maximum CPO drops from 3.87 to 2.83 W, in agreement with the trend observed by Jakovljevic et al. and Nathania et al. [4].

**Figure 3:**
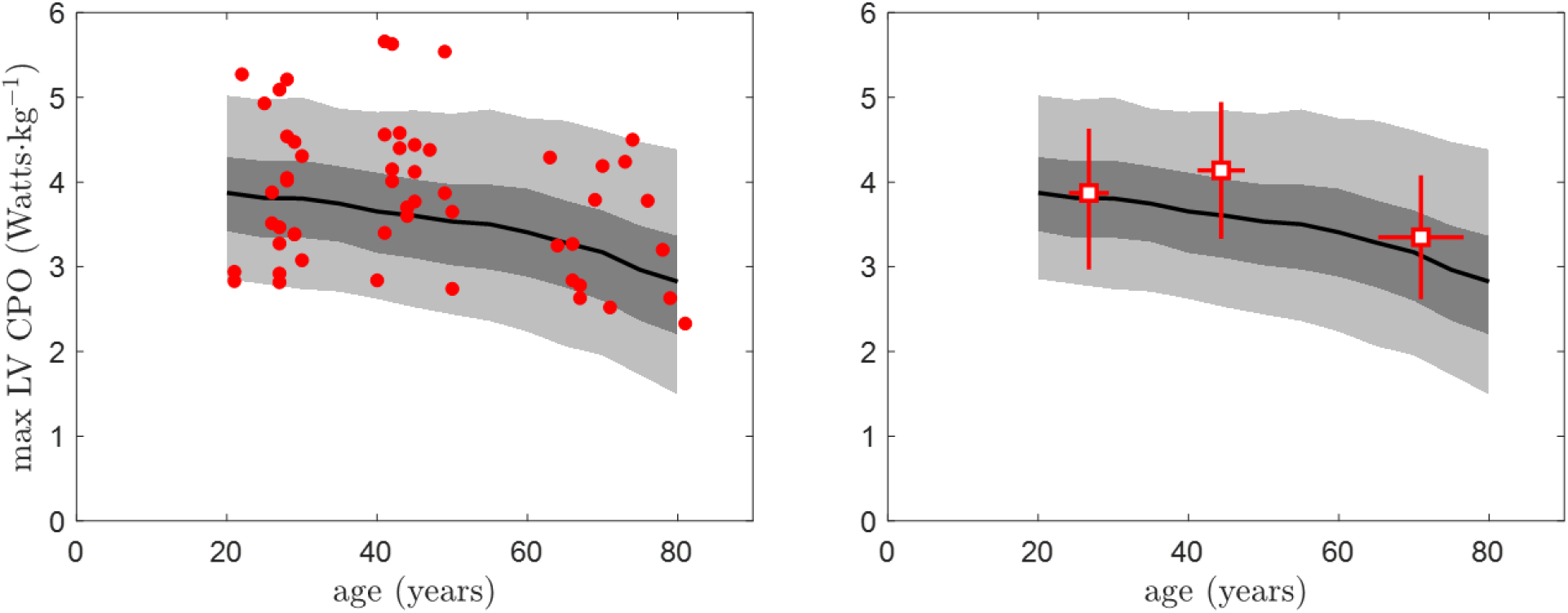
Predicted maximum left ventricular CPO as function of age. Both panels plot the population model predictions of maximum left-ventricular power output as functions age. The left panel shows observations from Jakovljevic et al. [3], with data from individual subjects plotted as individual data points. The right panel plots the means (open squares) and standard deviations (red lines) for the three age groups observed by Jakovljevic et al.

Similar to the resting state simulations in Figure 2, the majority of the data in Figure 3 falls within the 90% confidence range, with 8 of the 55 individuals falling outside of the 90% range and 20 of 55 data points falling within the 50% confidence range. The population model predicts an average drop of 0.48 W in maximum CPO from the middle age group (mean 44 years) to old group (mean 71 years), compared to an observed difference of 0.79 W. Maximum CPO falls below 2.2 W for roughly 25% of the simulated 80-year-old population, close to the value of 1.96 W that has been identified as the cut-off associated with maximum relative risk ratio for mortality [1]. Maximum CPO falls below 1.96 W for 15.5% of the simulated 80-year-old population.

## Discussion

Using an age-structured population model, the present study tested hypotheses that reductions in key cytosolic metabolite pools that occur with aging impair the myocardial capacity to synthesize ATP at physiological free energy levels, and that the resulting metabolic changes impair the mechanical pumping ability of the heart and its performance. Results support these hypotheses, as the model parameterized based on resting-state data predicts the observed relationships between age and mechanical and energetic performance of the heart in exercise. Specifically, the population-level model of human cardiac energetics was parametrized to match data on resting-state energetics in healthy women aged 21 to 81. Model simulations predict that metabolic supply capacity decreases with age, driven by changes in metabolic pools. Applying the assumption that maximum cardiac power output is limited by maximum metabolic supply yields predictions that are consistent with available data on reductions in maximum CPO with age, and are consistent with the following specific hypotheses:

1. reductions in cytoplasmic adenine nucleotide, creatine, and phosphate pools that occur with aging impair the myocardial capacity to synthesize ATP at physiological free energy levels; and
2. the resulting changes to myocardial energetic status play a causal role in contributing to reductions in maximal cardiac power output with aging.

The potential importance of the link between energetic status and mechanical function is highlighted by the fact that maximum CPO is the strongest predictor of mortality in heart failure [1]. Indeed, the hypotheses for mechanisms underlying age-dependent changes to myocardial mechano-energetic function tested here were formulated based on previous theoretical analysis of the link between energy metabolite levels and mechanical function of the heart in cardiac decomposition and failure [6, 7]. While these results do not suggest that the phenotypes are identical, they do suggest fundamental similarities in terms of mechanisms impeding myocardial energetics and mechanical-energetic coupling.

## Author Contributions

XG and DAB designed the study and conducted the analysis. All authors contributed to interpreting the results and writing the manuscript.

## Acknowledgments

This study supported by NIH grants U01HL122199 and HL144657. D.G.J. is a recipient of research funding from the Research Councils UK Newcastle Centre for Ageing and Vitality (Grant Number L016354) and the EU Horizon 2020 research and innovation programme under Grant Agreement Number 777204.

## References

1. Williams, S.G., et al., Peak exercise cardiac power output; a direct indicator of cardiac function strongly predictive of prognosis in chronic heart failure. Eur Heart J, 2001. 22(16): p. 1496–503.

2. Goldspink, D.F., et al., A study of presbycardia, with gender differences favoring ageing women. Int J Cardiol, 2009. 137(3): p. 236–45.

3. Jakovljevic, D.G., et al., Effect of physical activity on age-related changes in cardiac function and performance in women. Circ Cardiovasc Imaging, 2015. 8(1).

4. Nathania, M., et al., Impact of age on the association between cardiac high-energy phosphate metabolism and cardiac power in women. Heart, 2018. 104(2): p. 111–118.

5. Wu, F., J. Zhang, and D.A. Beard, Experimentally observed phenomena on cardiac energetics in heart failure emerge from simulations of cardiac metabolism. Proc Natl Acad Sci U S A, 2009. 106(17): p. 7143–8.

6. Tewari, S.G., et al., Dynamics of cross-bridge cycling, ATP hydrolysis, force generation, and deformation in cardiac muscle. J Mol Cell Cardiol, 2016. 96: p. 11–25.

7. Tewari, S.G., et al., Influence of metabolic dysfunction on cardiac mechanics in decompensated hypertrophy and heart failure. J Mol Cell Cardiol, 2016. 94: p. 162–175.

8. Ingwall, J.S., Energy metabolism in heart failure and remodelling. Cardiovasc Res, 2009. 81(3): p. 412–9.

9. Ingwall, J.S., On the control of metabolic remodeling in mitochondria of the failing heart. Circ Heart Fail, 2009. 2(4): p. 275–7.

10. Ingwall, J.S. and R.G. Weiss, Is the failing heart energy starved? On using chemical energy to support cardiac function. Circ Res, 2004. 95(2): p. 135–45.

11. Nascimben, L., et al., Creatine kinase system in failing and nonfailing human myocardium. Circulation, 1996. 94(8): p. 1894–901.

12. Neubauer, S., et al., Myocardial phosphocreatine-to-ATP ratio is a predictor of mortality in patients with dilated cardiomyopathy. Circulation, 1997. 96(7): p. 2190–6.

13. Ventura-Clapier, R., et al., Bioenergetics of the failing heart. Biochim Biophys Acta, 2011. 1813(7): p. 1360–72.

14. Stanley, W.C., Myocardial energy metabolism in dilated cardiomyopathy. Heart Metab., 2011. 49(5–8).

15. Mann, D.L. and M.R. Bristow, Mechanisms and models in heart failure: the biomechanical model and beyond. Circulation, 2005. 111(21): p. 2837–49.

16. Neubauer, S., The failing heart--an engine out of fuel. N Engl J Med, 2007. 356(11): p. 1140–51.

17. Bakermans, A.J., et al., Human Cardiac (31)P-MR Spectroscopy at 3 Tesla Cannot Detect Failing Myocardial Energy Homeostasis during Exercise. Front Physiol, 2017. 8: p. 939.

18. Ardehali, H., et al., Targeting myocardial substrate metabolism in heart failure: potential for new therapies. Eur J Heart Fail, 2012. 14(2): p. 120–9.

19. Gopal, D.M. and F. Sam, New and emerging biomarkers in left ventricular systolic dysfunction--insight into dilated cardiomyopathy. J Cardiovasc Transl Res, 2013. 6(4): p. 516–27.

20. Doenst, T., T.D. Nguyen, and E.D. Abel, Cardiac metabolism in heart failure: implications beyond ATP production. Circ Res, 2013. 113(6): p. 709–24.

21. Ashrafian, H., M.P. Frenneaux, and L.H. Opie, Metabolic mechanisms in heart failure. Circulation, 2007. 116(4): p. 434–48.

22. Kostler, H., et al., Age and gender dependence of human cardiac phosphorus metabolites determined by SLOOP 31P MR spectroscopy. Magn Reson Med, 2006. 56(4): p. 907–11.

23. Wu, F., et al., Phosphate metabolite concentrations and ATP hydrolysis potential in normal and ischaemic hearts. J Physiol, 2008. 586(17): p. 4193–208.

24. Olivetti, G., et al., Gender differences and aging: effects on the human heart. J Am Coll Cardiol, 1995. 26(4): p. 1068–79.

25. Ridout, S.J., et al., Age and sex influence the balance between maximal cardiac output and peripheral vascular reserve. J Appl Physiol (1985), 2010. 108(3): p. 483–9.

26. Tune, J.D., M.W. Gorman, and E.O. Feigl, Matching coronary blood flow to myocardial oxygen consumption. J Appl Physiol (1985), 2004. 97(1): p. 404–15.

27. Kates, A.M., et al., Impact of aging on substrate metabolism by the human heart. J Am Coll Cardiol, 2003. 41(2): p. 293–9.

28. Beard, D.A. and M.J. Kushmerick, Strong inference for systems biology. PLoS Comput Biol, 2009. 5(8): p. e1000459.

29. Knaapen, P., et al., Myocardial energetics and efficiency: current status of the noninvasive approach. Circulation, 2007. 115(7): p. 918–27.

